# Conserved serotonergic background of experience-dependent challenge-responding in zebrafish (*Danio rerio*)

**DOI:** 10.1101/785352

**Authors:** Zoltán K Varga, Diána Pejtsik, László Biró, Áron Zsigmond, Máté Varga, Blanka Tóth, Vilmos Salamon, Tamás Annus, Éva Mikics, Manó Aliczki

## Abstract

Forming effective responses to threatening stimuli requires the adequate and coordinated emergence of stress-related internal states. Such ability depends on early-life experiences and, in connection, the adequate formation of neuromodulatory systems, particularly serotonergic signaling. Here, we assess the serotonergic background of experience-dependent behavioral responsiveness employing a zebrafish (*Danio rerio*) model. For the first time, we have characterized a period during the behavioral metamorphosis in which zebrafish are highly reactive to their environment. Absence of social stimuli during this phase established by isolated rearing fundamentally altered the behavioral phenotype of post-metamorphic zebrafish in a challenge-specific manner, partially due to a decline in responsiveness and an inability to develop stress-associated arousal state. In line with this, isolation differently affected whole-brain 5-HT signaling in resting and stress-induced conditions, an effect that was present at the level of the dorsal pallium and was negatively associated with responsiveness. Administration of the 5HT1AR partial agonist buspirone prevented the isolation-induced serotonin response to novelty in the forebrain and rescued stress-induced arousal along with challenge-induced behaviors, which altogether indicates a functional connection between these changes. In summary, there is a consistent negative association between behavioral responsiveness and serotonergic signaling in zebrafish, which is well recognizable through the modifying effects of developmental perturbation and pharmacological manipulations as well. Our results imply a conserved serotonergic mechanism that context-dependently modulates environmental reactivity and is highly sensitive to experiences acquired during a specific early-life time-window, a phenomenon that was previously only suggested in mammals.

**Significance statement:** The ability to respond to challenges is a fundamental factor in survival. We show that zebrafish that lack appropriate social stimuli in a sensitive developmental period show exacerbated alertness in non-stressful conditions while failing to react adequately to stressors. This shift is reflected inversely by central serotonergic signaling, a system that is implicated in numerous mental disorders in humans. Serotonergic changes in brain regions modulating responsivity and behavioral impairment were both prevented by the pharmacological blockade of serotonergic function. These results imply a serotonergic mechanism in zebrafish that transmits early-life experiences to the later phenotype by shaping stress-dependent behavioral reactivity, a phenomenon that was previously only suggested in mammals. Zebrafish provide new insights into early-life-dependent neuromodulation of behavioral stress-responses.

## Introduction

Coping with environmental threats is essential for the survival and well-being of animals. Contextually adequate emergence and expression of behavioral responses depend on previous experiences, particularly stimuli acquired during sensitive and plastic developmental periods (Chapman et al., 2004; Haller et al., 2014; Nederhof and Schmidt, 2012; Santarelli et al., 2014). While there are numerous plausible explanations regarding the function of this link, e.g. preparing the individual for later environmental challenges, the underlying mechanisms are poorly understood. Clinical and preclinical data suggest that the quality of development of neuromodulatory pathways, particularly the formation of the serotonergic system, affects later behavioral phenotype, hence it transmits the information of the early-life environment to adulthood (Booij et al., 2015; Fone and Porkess, 2008; Gross et al., 2002; Lehmann et al., 2003; Lukkes et al., 2009b; Raleigh et al., 1984). The diverse and context-dependent behavioral effects of serotonin suggest that these pathways, apart from direct modulation, are able to act as general modulators of behavior (Geyer, 1995; Jing et al., 2009; Lovett-Barron et al., 2017; Yokogawa et al., 2012), possibly through the modification of internal states, such as arousal and alertness. State alertness adjusts behavioral reactivity to current environmental demands, e.g. the presence of acute stressors, by fine-tuning sensory-motor functions, information processing or decision-making processes. Despite its fundamental impact on the expression of behavior, there are only a few studies assessing the neuromodulatory basis of general arousal in healthy animals, while the literature of experience-dependent alertness and associated challenge-responding is even sparser.

The zebrafish (*Danio rerio*) provides an attractive tool to investigate basic vertebrate neurobiology, due to the ease and accessibility of approaches to modify or screen physiological processes (Lieschke and Currie, 2007; MacRae and Peterson, 2015). Particularly, the employment of zebrafish has several advantages to study the neural basis of experience-dependent challenge-responding: firstly, their relatively simple behavioral repertoire is primarily driven by robust stress responses including hormonal and neuromodulatory actions (Egan and Kalueff, 2009); secondly, their central nervous system possesses homologous anatomical structures and neurotransmitters to that of mammals with highly conserved function but less complex organization (Panula et al., 2010); thirdly, they show significant behavioral plasticity indicated by a robust behavioral reorganization occurring during the late larval stage (Lau and Guo, 2011). The onset of such metamorphosis gives us the opportunity to assess the impact of experiences in potentially sensitive developmental periods on the adult challenge-responding phenotype.

In the current study we aimed to utilize the above described unique features of zebrafish and assess how early-life events affect later challenge-responding phenotypes through alterations of 5-HT signaling. First, we aimed to identify a sensitive developmental period that plays a prominent role in shaping adult behavior. We hypothesized that the behavioral metamorphosis at the late larval stage is characterized by environmental reactivity that may serve as a critical period for shaping the individual’s later behavioural phenotype. Identifying such a sensitive period, we subsequently subjected zebrafish to the acute presence and/or chronic absence of environmental stimuli, e.g. social isolation and analyzed different aspects of challenge responsiveness. In order to do this, we conducted tests that represent non-meaningful stimuli i.e. an unidentified threat, such as environmental novelty and meaningful stimuli, such as a social cue. These contexts evoke different internal states and behavioral responses that may be orchestrated by different neuromodulatory actions (reviewed by Ralf-Peter Behrendt, 2011). We also differentiated between novel visual, social visual and novel but non-visual stimuli in order to exclude potential biasing effects of social isolation on sensory perception. Finally, we assessed isolation-induced changes in serotonergic signaling under resting and challenge-induced conditions by applying molecular, histological and pharmacological tools. We conducted measurements at whole-brain level and more specifically in limbic forebrain areas, then functionally correlated serotonergic and behavioral effects by the pharmacological blockade of 5-HT signaling.

## Materials and Methods

### Animals

Wild type (AB) zebrafish lines were maintained in the animal facility of Eötvös Loránd University according to standard protocols (Westerfield, 2000). Experimental subjects were unsexed animals aged between 8 and 30 dpf (days post fertilization). Fish were maintained in a standard 14 h/10 h light/dark cycle. Animals were terminated on ice immediately after each experiment. Feeding of larvae started at 5 dpf with commercially available dry food (a 1:1 combination of < 100 μm and 100–200 μm Zebrafeed, Sparos) combined with *Paramecium*. After 15 dpf, juvenile fish were fed using dry food with gradually increasing particle size (200–400 μm Zebrafeed, Sparos) combined with fresh brine shrimp hatched in the facility. All protocols employed in our study were approved by the Hungarian National Food Chain Safety Office (Permits Number: PEI/001/1458-10/2015 and PE/EA/2483-6/2016).

### Environmental modifications

Chronic social isolation (SI) of zebrafish was carried out between 14 and 29 dpf. Isolated animals were kept in white opaque plastic tanks (52 x 35 x 46 mm, depth x width x length) (Figure 2a) depriving the individuals of sensory cues from conspecifics. Tanks were filled with E3 medium, half of the volume of which was replaced on a weekly basis. Socially reared control animals were subjected to similar conditions with the exception of the size of their aquarium, which was matched to the greater number of larvae, providing approximately 50 ml volume for each larva.

**Figure 1:**
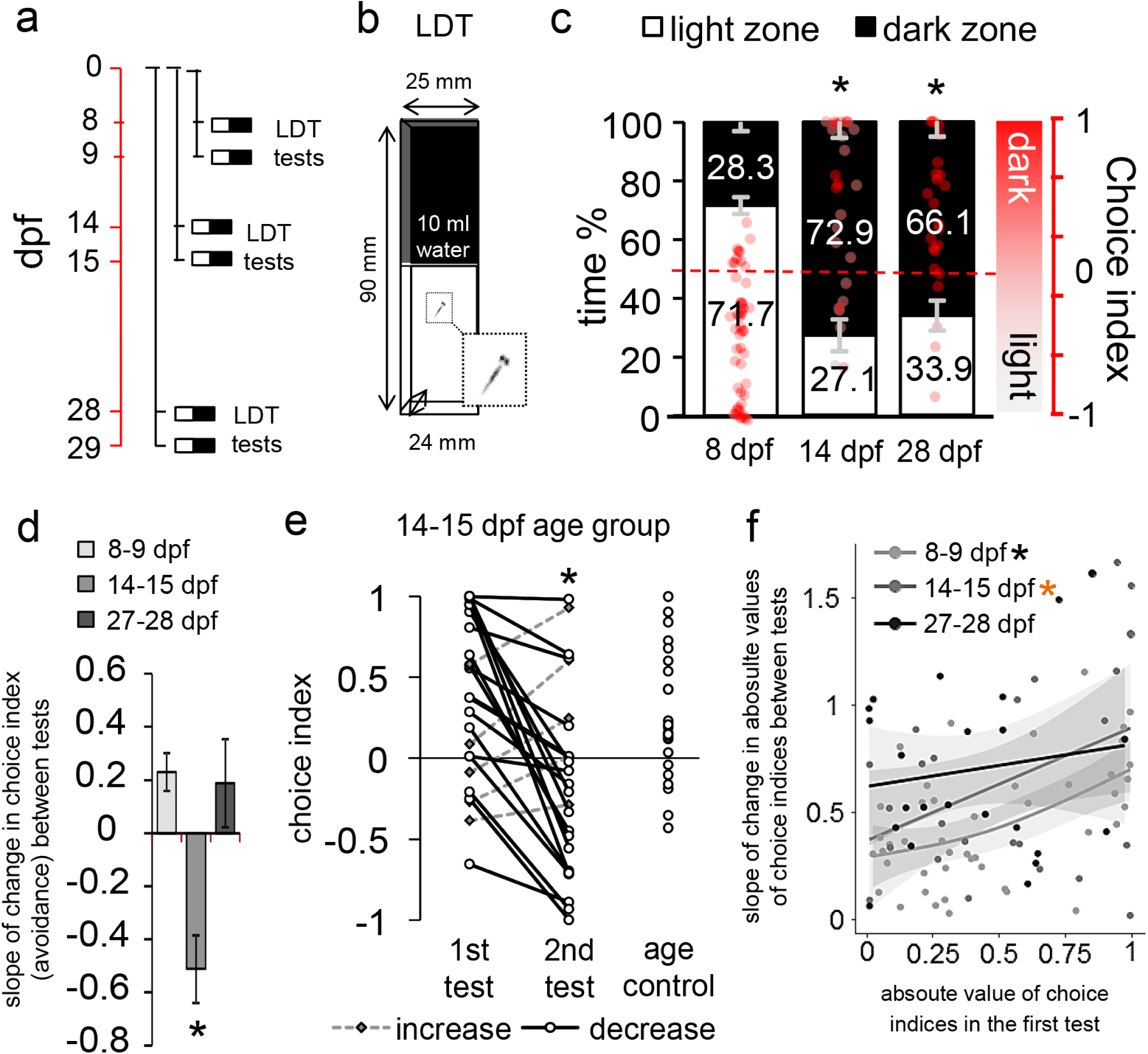
**a)** Protocol of *Experiment 1*. Red timeline shows the age of larvae expressed in days post fertilisation (dpf). Individual black lines show different groups of larvae. **b)** Schematic figure of the LDT apparatus. **c)** Avoidance behavior of zebrafish at the 1st testing day throughout the first month of development. Left axis shows the percentage of time, right axis shows choice index in which 1 indicates 100% time in the dark zone, whereas −1 represents 100% time in the light zone. Asterisk indicates significant difference compared to 8 dpf age group. **d)** Slope of change in avoidance behavior between the two testing days. Asterisk indicates significant difference compared to 8 dpf age group. **e)** Individual values of choice indices in the 14-15 dpf age group and test naïve age control. Dashed lines mark increased avoidance, solid lines indicate decreased avoidance. Asterisk indicates significant difference from 1st testing day. **f)** Relation between the first and second days of testing. Trends are indicated by solid regression lines and 95% confidence intervals. Black asterisk indicates significant linear regression. Orange asterisk indicates marginal significance (p=0.0583).

**Figure 2:**
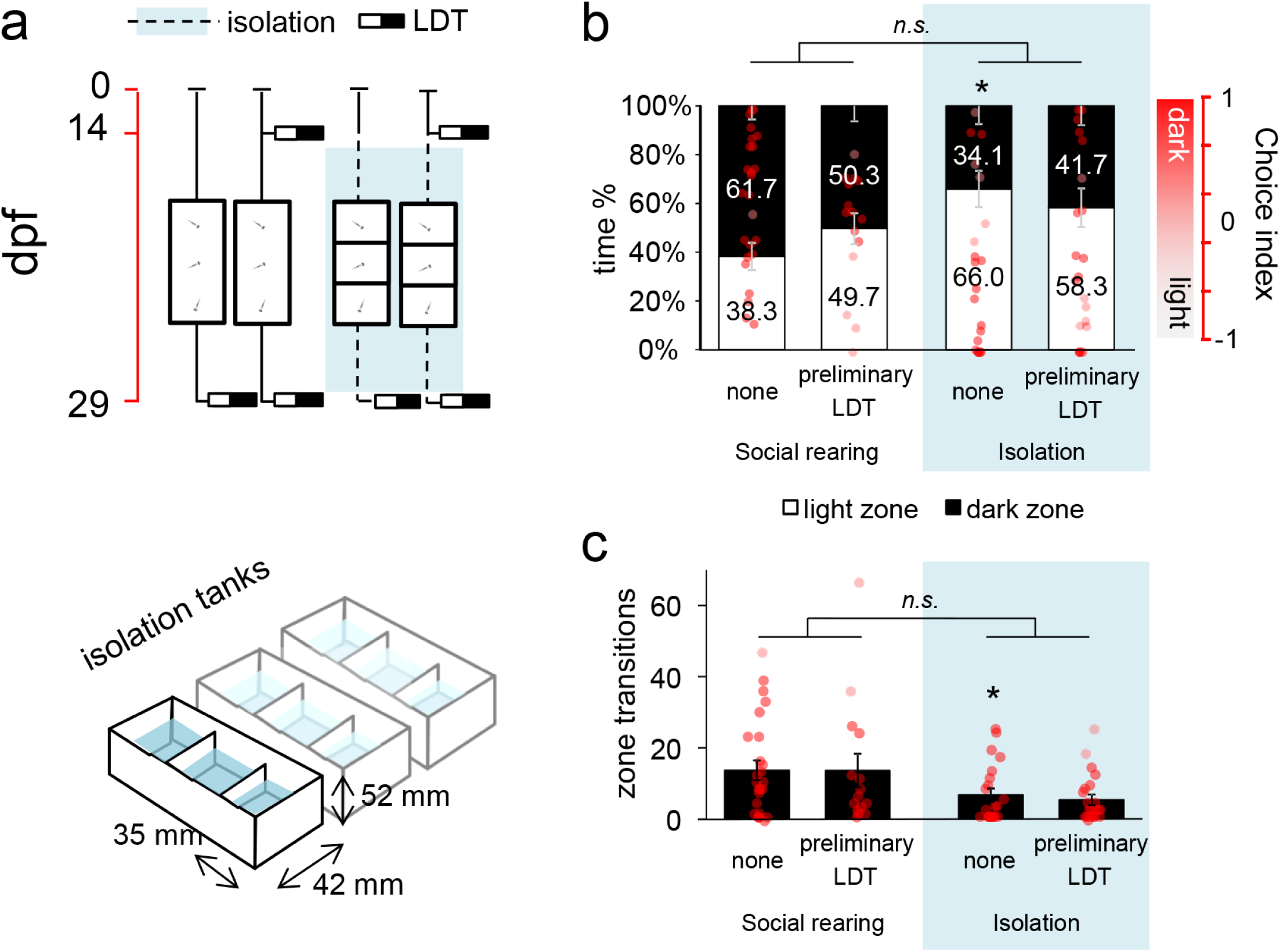
**a)** Protocol of *Experiment 2* (top) and schematic drawing of the isolation tanks (bottom). Red timeline shows the age of larvae expressed in days post fertilisation (dpf). **b)** Avoidance behavior of post-metamorphic zebrafish 2 weeks following a single LDT challenge and/or after 2 weeks of chronic social isolation. Left axis shows the percentage of time, right axis shows choice index in which 1 indicates 100% time in the dark zone, whereas −1 represents 100% time in the light zone. The reference levels of the linear mixed model were „social rearing” and „treatement none”. Asterisk indicates differences between rearing conditions at the level „treatment none”, n.s. indicates no significant differences between the relation of the treatment levels in socially reared and isolated groups. **c)** Number of transitions between light and dark zones of post-metamorphic zebrafish 2 weeks following a single LDT challenge and/or after 2 weeks of chronic social isolation.

Light/dark tank (LDT) exposure, considered as an acute challenge, was used as a developmental experience as well as a behavioral sampling tool. The procedure was identical in both cases and is described in the *Behavioral tests and analysis* section.

### Drug treatments

Buspirone (Sigma-Aldrich) was dissolved in E3 medium and administered as a water bath for 10 minutes followed by a 1-min washout in compartments of a 24-well plate. Each compartment (15.6 mm in diameter) contained 1.5 ml of treatment solution. We applied 0 (vehicle) and 50 mg/l concentrations which were selected based on our previous studies (Varga et al., 2018).

### Behavioral tests and analysis

For light/dark tank (LDT) testing, half of the base of a plastic tank (90 x 25 x 24, length x width x depth) (Fig1a) was masked with opaque matte black paint (dark zone), while the other half remained intact, penetrable by light (light zone). Experiments were conducted during the second part of the light phase of the daily cycle, as zebrafish are reported to show continuous activity in this period (MacPhail, 2009). 8 subjects were tested simultaneously over the span of a single trial. Experiments in which pharmacological treatments were employed began with a 10-minute treatment bath in a 24-well plate compartment after which animals were gently pipetted to another compartment containing E3 medium for a 1-minute washout and then were moved to the testing apparatus for 10 minutes. Behavior of fish was recorded by a digital video camera. Animals were placed in the light zone of the apparatus directly next to the border of zones. Experimental tanks (Figure 1b) were each filled with 10 ml medium and thoroughly cleaned between trials. To reduce interfering stimuli from the environment, experimental apparatuses were covered with an opaque black plastic box with a hole on top allowing the attachment of the video camera. The apparatuses were illuminated from beneath with white LED panels covered by matte Plexiglas. Video recordings were analyzed using EthoVision XT 12 automated tracker software (NOLDUS L.P.J.J. et al., 2001) and Solomon Coder (András Péter, 2017) manual tracker software. Time spent in and transitions between light and dark zones were measured. Distance swum and mean velocity were measured only in the light zone. To describe scotophobia (dark avoidance) and scototaxis (dark preference) a choice index was defined as the time spent in the dark zone of the compartment relative to time spent in the light zone. Consequently, choice index of 1 indicates 100% time in the dark zone, choice index of −1 represents 100% time in the light zone and choice index of 0 represents 50% in the dark and 50% in the light zone.

To measure arousal and visual responsiveness of fish, standard, transparent 6-well plates (34.8 mm in diameter) were used. Illumination and visual stimuli were presented by a screen of a tablet PC from beneath. The choice of time for testing and the applied pharmacological treatment protocols were the same as those in earlier experiments. The experiment consisted of two parts; one was considered as a resting and the other was considered as an acute stress-affected period. To establish basal conditions, a 20-min habituation phase was introduced before the first part, as zebrafish have been reported to habituate to novelty in this amount of time (Yokogawa et al., 2012). To measure behavior in stress-affected conditions the second part took place immediately after the transfer of animals to a novel compartment. In order to measure arousal state and accompanied sensory responsiveness of the animals, during both parts a black object was projected onto the bottom of the testing compartment that remained stationary for 5 minutes, then started to rotate in circles with its speed increasing 2-fold in every 2 minutes, from 0.5 rpm to 8 rpm, for an overall 10 minutes (see supplementary video 1). Both the initial position and subsequent movement directions of the items were randomized among test subjects. The number of active swimming episodes was measured with a custom-made clustering algorithm based on color clustering (k-means clustering). We considered a swimming episode active if more than 1% pixel change occurred between two consecutive frames. This threshold was calculated empirically from consecutive frames showing stationary or moving animals.

To measure avoidance responses to non-meaningful, e.g. novelty-induced, non-visual challenges, the swimming plus-maze (SPM) test was used, recently developed by our laboratory (Varga et al., 2018). The apparatus is a “+” shaped platform consisting of two plus two opposite arms, different in depth, connected by a center zone. Larval and juvenile zebrafish prefer the deep over the shallow arms and the center zone, a preference linked to anxiety-like motivational states. For the validation procedure and detailed specification of the test see Varga et al. (2018). 10 minutes long behavioral tests were conducted employing the same protocol as described at the LDT. Time spent in each zone and the mean of overall velocity were measured. To characterize arm preference, a choice index was defined as the relative time spent in the deep compared to the time spent in the shallow arms. Consequently, a choice index of 1 indicates 100% time in the deep arms, whereas a choice index of −1 represents 100% time in the more aversive shallow arms and a choice index of 0 represents an equal amount of time in each arm.

To measure avoidance responses to meaningful, e.g. social visual challenges the social preference test of the Dreosti laboratory was used (Dreosti et al., 2015). The test apparatus is a U-shaped platform (40 x 32 mm) consisting of two identical arms, with glass window partitions enabling only visual communication between zebrafish. The test is based on the visually-driven preference of larvae towards conspecifics, a phenomenon emerging around the third week of development. 15-minute-long behavioral tests with the presence of a conspecific were preceded by a 15-minute-long habituation phase according to the protocol from the original paper. Time spent in each zone was measured. To characterize social preference, a choice index was defined as the relative time spent in the arm with the conspecific compared to the time spent in the empty arm. Consequently, a choice index of 1 indicates 100% social preference, a choice index of −1 represents 100% social aversion and a choice index of 0 represents no preference or aversion.

### High Performance Liquid Chromatography-tandem mass spectrometry (HPLC/MS/MS)

Zebrafish were terminated and kept at −20 ◦C until measurements. The body and eyes of each subject were manually removed on ice cold petri dishes, then tissue samples were pooled from the remains of two randomly chosen individuals of the same treatment group. The samples were transferred to Eppendorf tubes and their weight was measured on a semi-micro balance (Precisa EP 225SM-DR). After the addition of 50 µl of 0.1% formic acid solution containing 5% methanol, the samples were homogenized using an ultrasonic sonotrode, and the homogenates were centrifuged to produce protein-free supernatants for HPLC-MS-MS analysis. The separation was carried out by high performance liquid chromatography composed of a Perkin Elmer series 200 high pressure gradient pump, an autosampler, an online degasser and a thermostat. A Phenomenex Synergi Hydro-RP 80 150*3,0 mm, 4 μm column was used with a gradient elution with methanol (mobile phase A) and 0.1% formic acid (mobile phase B): 0 min to 1.5 min: at 5% A; 3 to 4 min: at 60% A; 4.5 to 7 min: at 5% A. 10 µl samples were injected, and the flow rate was 500 µl/min. Serotonin was detected using a triple quadrupole mass spectrometer (Applied Biosystems MDS SCIEX 4000 Q TRAP) in positive multiple reaction monitoring mode (MRM transition: 177.2 → 170.0).

### Immunohistochemisty and confocal microscopy

Zebrafish were fixed overnight in 4% paraformaldehyde in phosphate-buffered saline (PBS). For the preparation of cryosections, cryoprotecting dehidration of whole-body samples was carried out in a 10-20-30% sucrose-PBS gradient. Samples were subsequently embedded in Tissue-Tek OCT medium, frozen on dry-ice and stored at −80°C until cryosectioning. 20-µm sections were cut in a Microm HM505 cryostat. Sections were collected onto slides (Thermo Fisher), dried at room-temperature overnight, and stored at −80°C until immunohistochemistry was performed.

For immunohistochemistry, slides with sections were bordered with liquid blocker using a 2mm wide PAP pen (Merck). For each step, 400 µl liquid solution was administered for each slide. Sections were washed three times in PBS (10 min each) and incubated overnight at room temperature with polyclonal anti-5-HT antibody produced in rabbit (1:500, 20080, ImmunoStar) diluted in PBS containing 0.5% bovine serum albumin (A7906, Sigma-Aldrich, Hungary) and 0.25% Triton X-100 (T8787, Sigma-Aldrich, Hungary). Sections were washed three times in PBS and incubated with goat anti-rabbit Alexa Fluor 488 (1:500, 111-545-003, Jackson ImmunoResearch Laboratories Inc.) for 2.5 hours. After several washes in PBS, sections were incubated with DAPI (1:10000, D8417, Sigma-Aldrich, Hungary) for 30 min. Slides were mounted with Mowiol4–88 (81381, Sigma-Aldrich, Hungary) fluorescent mounting medium.

Images were taken using a C2 confocal laser scanning microscope (Nikon Europe, Amsterdam, The Netherlands) using a 20× (Plan Apo VC, NA= 0.75) objective (xy: 0.31μm/pixel). We used 405 (Coherent) and 488 (CVI Melles Griot) lasers, and scanning was done in channel series mode. The pixel dwell time, PMT gain, laser intensity detector, and pinhole settings were kept constant for all image acquisitions.

Image analysis was carried out using the ImageJ software (Schneider et al., 2012). Standard 30 x 30 µm regions of interest (ROI) were chosen and applied in each investigated area.

Individual subtract values of a given image were calculated in a neighboring ROI, containing no fibers, by the enhancement of the subtraction to a level in which the background was totally diminished. Subsequently, chosen values were applied to relevant ROIs and the percentage of 5-HT signal in a binary processed picture was measured. Individual data were calculated as means of bilateral measurements.

### Experimental design

In *Experiment 1*, we aimed to determine the time point of shift from the light to the dark preferring phenotype during development, representing a specific aspect of the behavioral metamorphosis, by subjecting different sets of zebrafish to the LDT test for 10 minutes at 8, 14 and 28 dpf, respectively. Sampling times were chosen according to the measures of Lau and Guo (2011). To analyze environmental reactivity as well in these developmental stages, we repeated the tests on the subsequent days with the same animals and analyzed habituation capacity (Figure 1a). Different sets of larvae from the same spawning were tested in each 2-day trial set. Between the two tests animals were kept in the isolation tanks described before. Samples sizes were 48 for the 8 dpf, 24 for the 14 dpf and 23 for the 28 dpf age groups.

In *Experiment 2*, we investigated whether chronic absence of social stimuli affects post-metamorphic juvenile behavior and reactivity to acute environmental stimuli (LDT test). We subjected 14 dpf zebrafish to the LDT test and/or 2 weeks of social isolation, then analyzed their avoidance behavior in the LDT test (Figure 2a) at 29 dpf. Sample sizes were 25 and 21 (naïve and test-experienced) for the socially reared and 15 and 21 (naïve and test-experienced) for the isolated groups.

In *Experiment 3*, we aimed to determine isolation-induced changes in central nervous system 5-HT content under basal and challenge-induced conditions by conducting HPLC-MS measurements on juveniles before, immediately after and 1.5 hours after LDT testing. Sample sizes were 22, 18 and 24 (basal, immediate and 1.5 hours) for the socially reared and 16 for the isolated groups.

In *Experiment 4a*, in order to localize isolation-induced changes in 5-HT responses to acute challenges, we subjected juveniles to the LDT test and analyzed 5-HT immunoreactivity throughout the telencephalon 1.5 hours after the stress exposure. To unravel the connection between the observed behavioral and area-specific physiological changes, we aimed to diminish 5-HT activity by the administration of the anxiolytic agent buspirone, predominantly acting on 5-HT1A autoreceptors (Figure 4a). Sample sizes were 31 and 22 (vehicle and buspirone-treated) for the socially reared and 14 and 16 (vehicle and buspirone treated) for the socially isolated groups.

In *Experiment 4b*, we aimed to investigate the effects of isolation under basal and stress-induced arousal states and accompanying sensory responsiveness by subjecting juveniles to stationary and moving items projected onto the testing arena. To clarify the role of the serotonin response we treated the animals with buspirone between basal and stress-induced phases. Sample sizes were 13 and 14 (vehicle and buspironetreated) in both socially reared and isolated groups as well.

In *Experiment 4c* and *4d*, we aimed to investigate the effects of isolation in different contexts by subjecting juveniles to the SPM or the U-shaped social preference tests after a single treatment of buspirone. In *Experiment 4c*, sample sizes were 28 and 24 (vehicle and buspirone treated) for the socially reared and 17 and 16 (vehicle and buspirone treated) for the socially isolated groups, while in *Experiment 4d* these were 16 and 25 (vehicle and buspirone-treated) for socially reared and 14 for each isolated group.

### Data analysis

Data are shown as mean ± SEM. We performed statistical analyses in R Statistical Environment (R Core Team, 2017). To analyze the interactional effects of rearing conditions, developmental experiences and pharmacological treatments we used linear mixed models (Julian J. Faraway, 2016) from the *lme4* package (Douglas Bates, 2015). Rearing “social” and treatment “vehicle” were set as reference levels. To separate variance stemming from time or sequence of experimental trials or location of test platforms, these factors were added as random effects to our models. The statistical output of such models indicate if the effects of treatments are identical throughout rearing conditions. In this manner, we only showed the results in relation to the reference category, e.g. the difference between control and treated isolated groups differs from the corresponding difference between socially reared groups. To analyze within-group differences between repeated measures, we fitted linear mixed models with subject identifiers as random effects. We computed Pearson correlation coefficients (*r*) using the *GGally* package (Barret Schloerke et al., 2017). We calculated *p*-values from *t*-values of *lme4* using the *lmerTest* package (Kuznetsova et al., 2016) and rejected H_0_ if *p*-values were lower than 0.05.

## Results

### Behavioral metamorphosis in zebrafish defines a time-window with enhanced environmental reactivity

Since larval and juvenile zebrafish express a different behavioral repertoire (Dreosti et al., 2015; Lau and Guo, 2011) it is evident that, besides the morphological metamorphosis, a behavioral reorganization also occurs in this species. We hypothesized that the time period of such transformation may also function as a window of opportunity for individuals to sample current environmental demands and adapt to potential challenges. Such processes require a great amount of environmental reactivity and may represent a sensitive developmental period that may be suitable to model the effects of early-life experiences. In *Experiment 1* (Figure 1), we aimed to determine the time of the switch from the light to the dark preferring phenotype, representing a specific aspect of the behavioral metamorphosis (Lau and Guo, 2011), by subjecting separate sets of zebrafish to the LDT test for 10 minutes at 8, 14 and 28 dpf respectively (Figure 1a). To characterize environmental reactivity as well in these phases of development, we repeated the experiments the following day (9, 15, or 29 dpf respectively) with the same animals. While 8 dpf larvae preferred the light over the dark zone, the preference was shifted towards the dark zone in older groups, (Figure 1c) outlining the time onset of the aforementioned phenotypic shift. At test repetitions, 9 dpf light-preferring larvae showed increased dark exploration, 15 dpf dark preferring larvae showed increased light exploration, while 29 dpf fish expressed no detectable change of behavior compared to the first testing day (Figure 1d). It is important to note that 15 dpf larvae showed the most extreme inter-test decrease indicated by the slope of change in avoidance (Figure 1d), and this group had the highest proportion of animals showing habituation, 19 out of 24 fish decreased their avoidance (Figure 1e). Furthermore, we did not detect any differences between 14 and 15 dpf test-naïve animals, indicating that the observed effects were experience-dependent changes and were not biased by aging. Linear regression showed that in the 8 and 14 dpf groups, the absolute value of avoidance in the first tests predicted the absolute value of slope of change in avoidance between tests (Figure 1f). Extreme first-trial avoidance was associated with stronger inter-test changes in behavior. In summary, the first 2 weeks of development mark a specific aspect of behavioral metamorphosis and a period of enhanced reactivity that peaks at the second sampling event. See statistical data in Table 1.

**Table 1:**
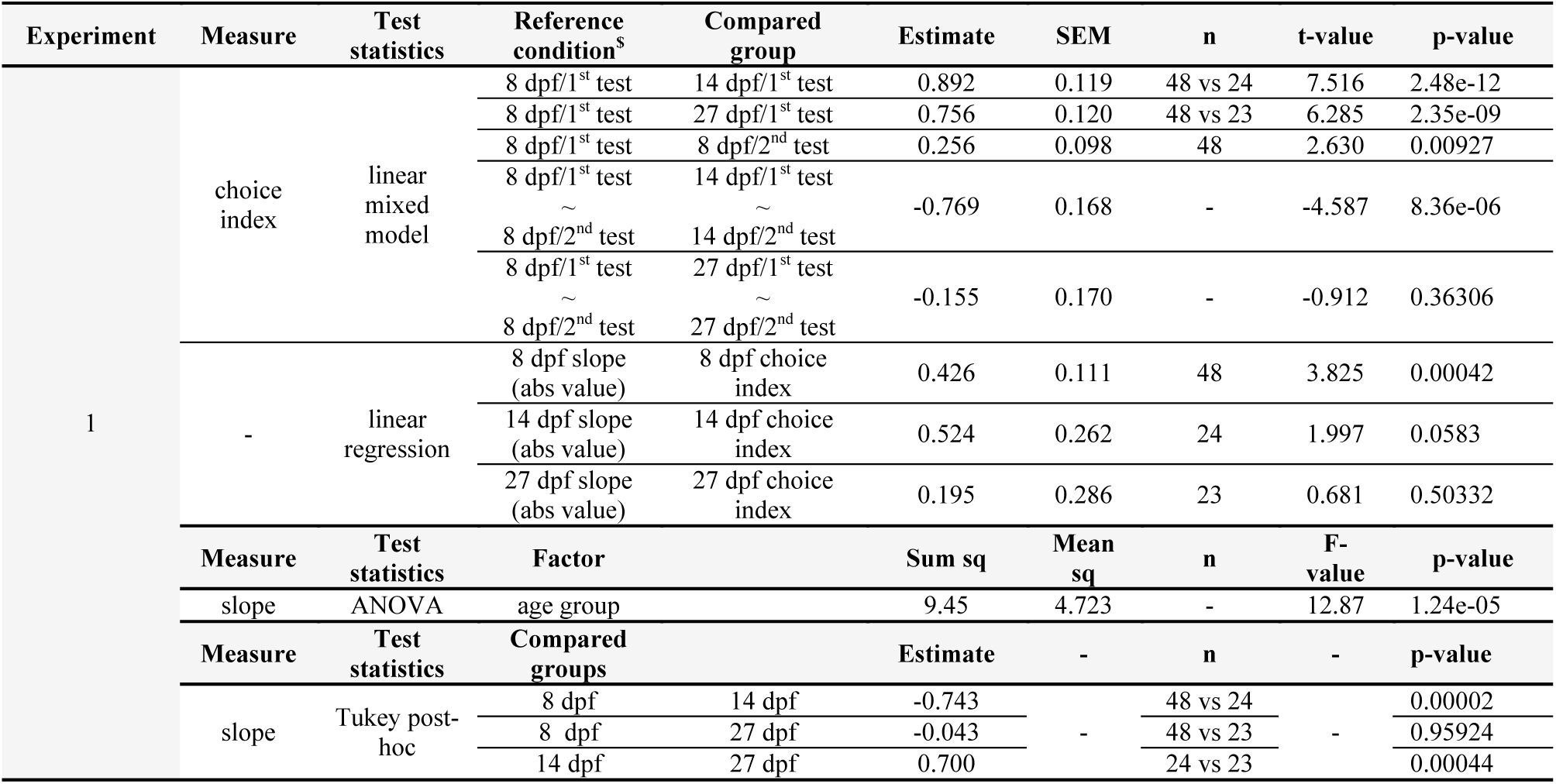
Statistical data of *Experiment 1*. ^$^ The statistical output of these linear mixed models are able to show significant differences from a reference condition in each analyzed levels, i.e. age and testing day. Age “8 dpf” and testing day “1^st^” were set as reference conditions. We also show interactional effects in relation to the reference categories, e.g. the difference between the 1^st^ and 2^nd^ tests (∼) at 8 dpf differ from the corresponding difference between the 14 dpf age groups. Note that different hypotheses testing methods were used, hence shown parameters may differ from method to method.

### Social isolation in the reactive period disrupts later behavioral responsiveness of zebrafish

Given such increased environmental reactivity around the second week of development, in *Experiment 2* we assessed the effects of different experiences gained in this time-window on the post-metamorphic behavioral phenotype. We subjected zebrafish larvae to acute novelty stress (LDT testing) at 14 and/or chronic environmental isolation from 14 to 28 dpf, then analyzed their avoidance behavior in the LDT test at the juvenile stage (Figure 2a). Note that the chronic environmental deprivation started at the most plastic sampling point where behavioral development has not occurred yet and ended after the morphological metamorphosis, hence we excluded the chance of detecting a possible conservation event of the immature avoidance phenotype. Environmental isolation, regardless of the onset of preliminary LDT testing, decreased avoidance behavior (Figure 2b), the number of transitions between zones (Figure 2d) and mean velocity (data not shown), indicating an overall impairment in responsivity. See statistical data in Table 2.

**Table 2:**
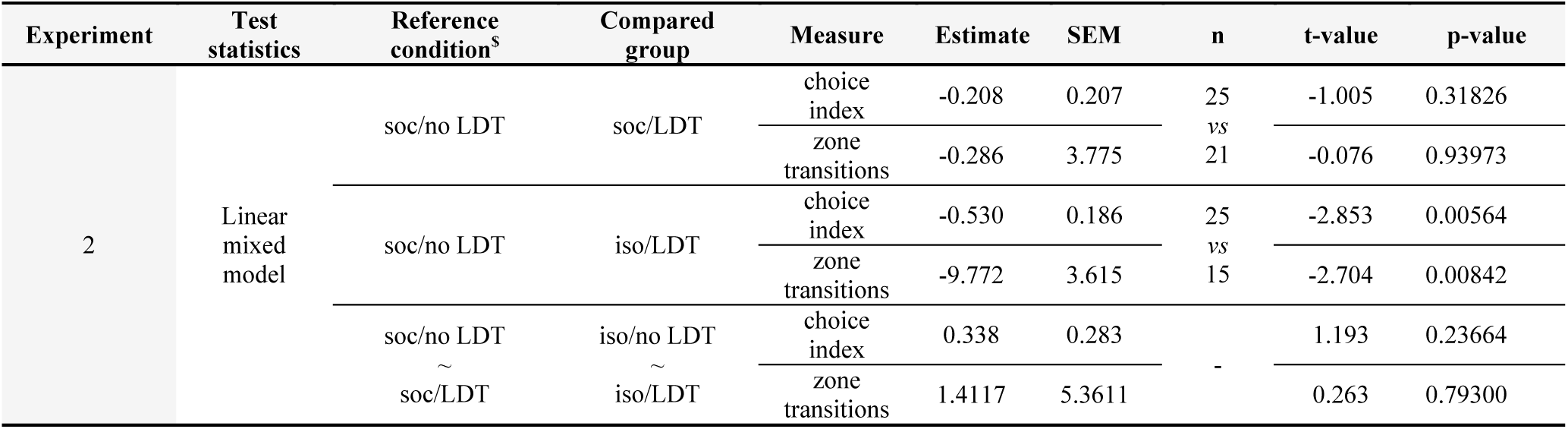
Statistical data of *Experiment 2*. ^$^ The statistical output of these linear mixed models are able to show significant differences from a reference condition in each analyzed levels, i.e. rearing condition and prior testing. Rearing “social” and prior testing “no LDT” were set as reference conditions. We also show interactional effects in relation to the reference categories, e.g. the difference between the “no LDT” and “LDT” groups (∼) in socially reared groups do not differ from the corresponding difference between the isolated groups.

### Social isolation differentially affects central 5-HT levels of zebrafish under resting and novel conditions

The decreased response capacity observed led us to investigate the role of 5-HT in isolation-induced effects, since serotonergic signaling is linked to behavioral constraint and environmental responsiveness as well (for reviews see Brodie and Shore, 1957; Depue and Spoont, 1986; Lucki, 1998; Soubrié, 1986; Spoont, 1992). To characterize the dynamics of potential 5-HT level changes in our isolation model, we conducted HPLC-MS-MS measurements on whole-brain samples of separate sets of zebrafish under resting conditions, as well as immediately- and 1.5 h after the post-metamorphic LDT test, respectively (Figure 3a). Socially reared fish showed a rapid decrease in 5-HT content in response to LDT testing, changes that were detectable for at least as long as 1.5 hours. In contrast, isolated fish showed decreased baseline levels compared to socially reared conspecifics and a markedly enhanced, transient 5-HT response to LDT testing (Figure 3b). Such differentially directed 5-HT responses to the novelty-challenge were accompanied by a similar decrease in light avoidance that we measured in *Experiment 2*. Interestingly, minute-by-minute analysis revealed that the principal component of such low responsivity was the delayed emergence of avoidance activity, while decreased locomotion was stable throughout the test (Figure 3c and 3d). See statistical data in Table 3.

**Figure 3:**
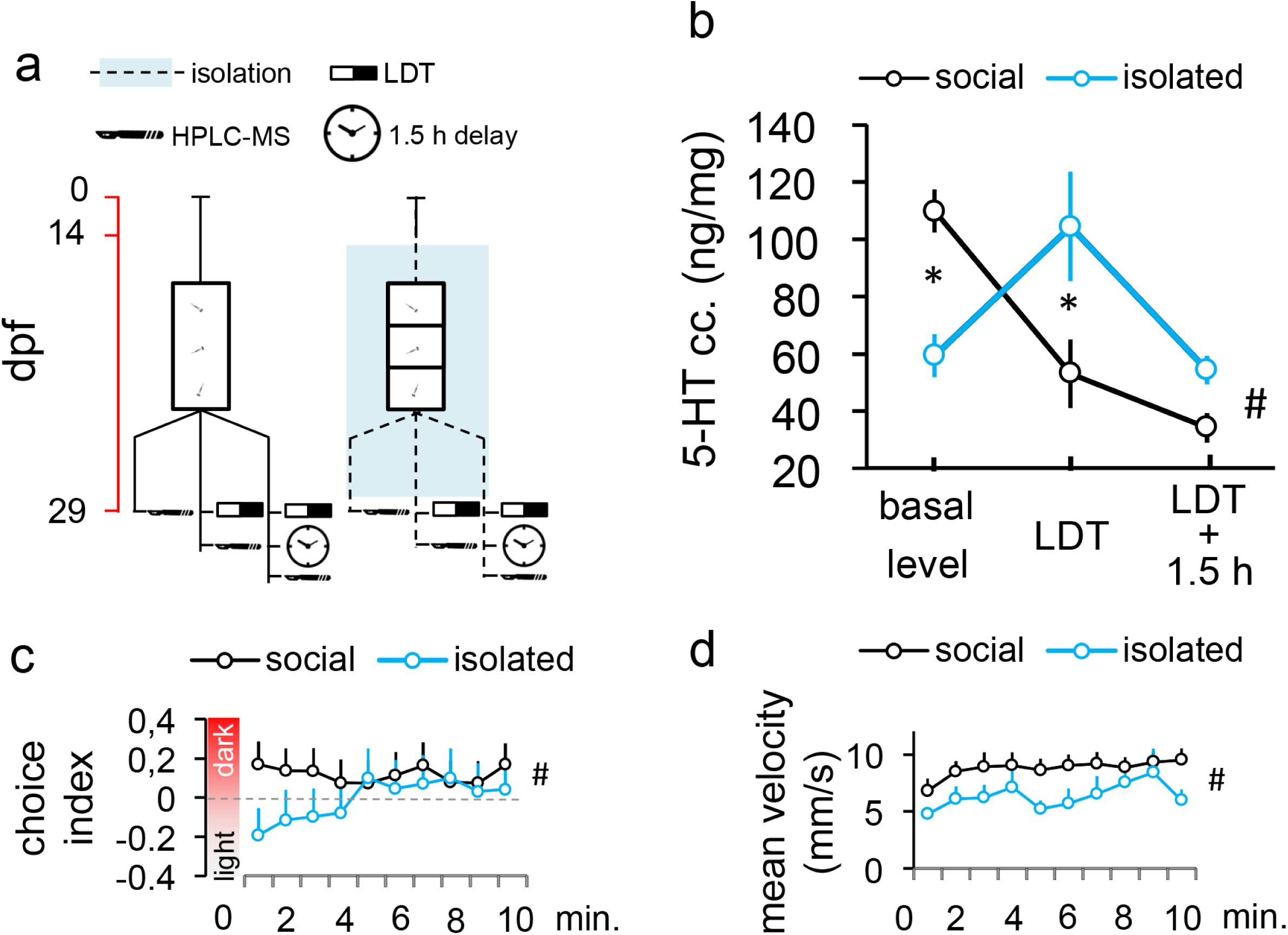
**a)** Protocol of *Experiment 3*. Red timeline shows the age of larvae expressed in days post fertilisation (dpf). **b)** Whole-brain 5-HT concentration of post-metamorphic zebrafish in resting conditions, and immediately and 1.5 hours after LDT challege. Asterisk indicates differences between groups at the level of a single condition (basal level, LDT, LDT+1.5 h), hashtag indicates significant group*condition interaction. **c)** Minute-by-minute resolution analysis of avoidance measured by choice index and **d)** velocity. Hashtag indicates significant group*time interaction.

**Figure 4:**
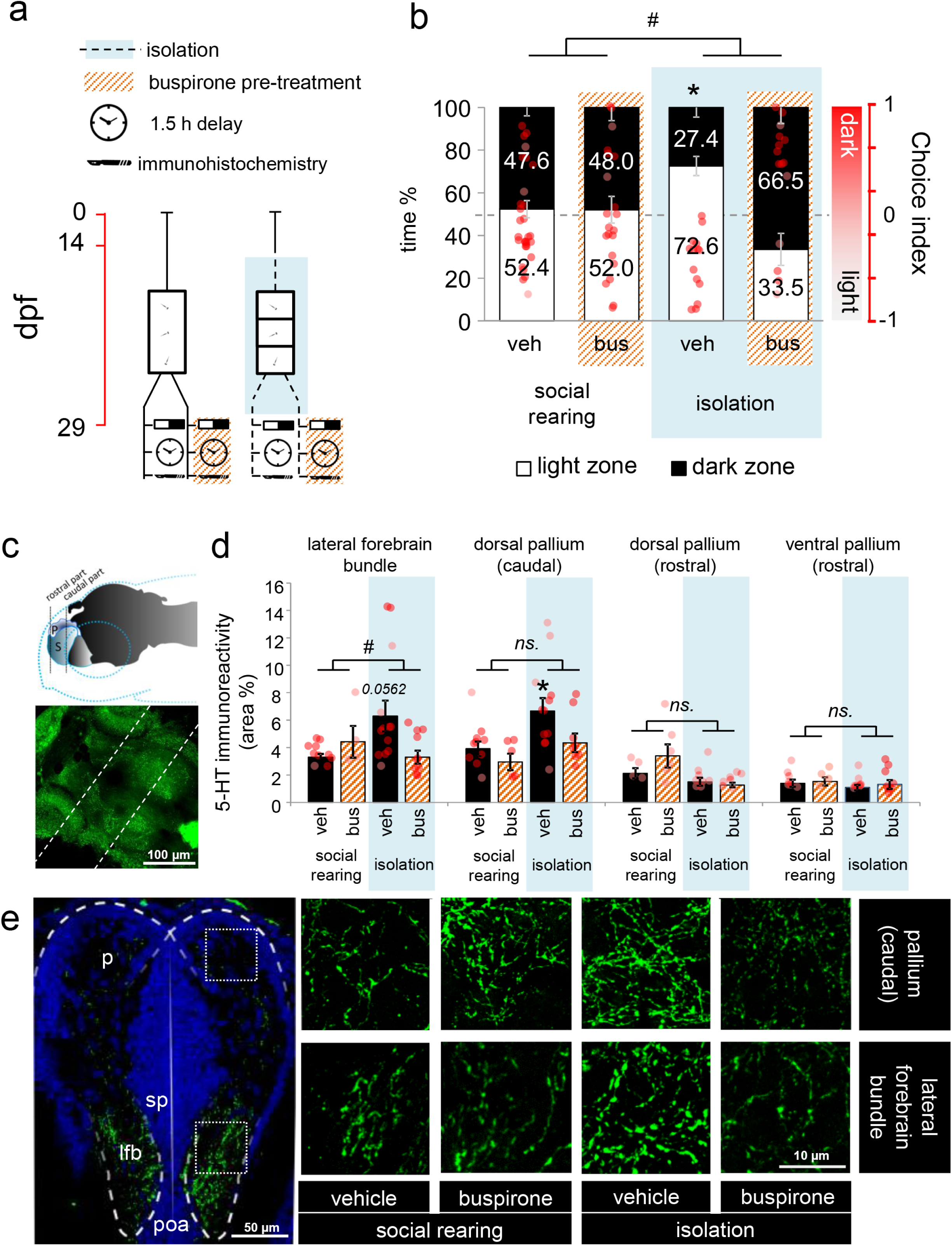
**a)** Protocol of *Experiment 4a*. **b)** Avoidance behaviour of post-metamorphic zebrafish 2 weeks following chronic social isolation and/or after acute buspirone treatment. Left axis shows the percentage of time, right axis shows choice index in which 1 indicates 100% time in the dark zone, whereas −1 represents 100% time in the light zone. Reference levels of the linear mixed model were „social rearing” and „treatement vehicle”. Asterisk indicates differences between rearing conditions at the level of treatment vehicle, hashtag indicates significant differences between the relation of the treatment levels in socially reared and isolated groups. **c)** Schematic drawing (top) and confocal microphotographs (maximum intensity projection) (bottom) of the investigated areas. The image was taken from a whole-brain preparatum of a zebrafish. Dashed lines indicate the approximate anterio-posterior position of the investigated areas. **d)** Quantification of 5-HT immunoreactivity in the analized brain regions. Asterisk indicates differences between rearing conditions at the level of treatment vehicle, hashtag indicates significant differences between the relation of the treatment levels in socially reared and isolated groups. **e)** Representative confocal microphotographs (maximum intensity projection) of sites with significant differences in immunoreactivity.

**Table 3:**
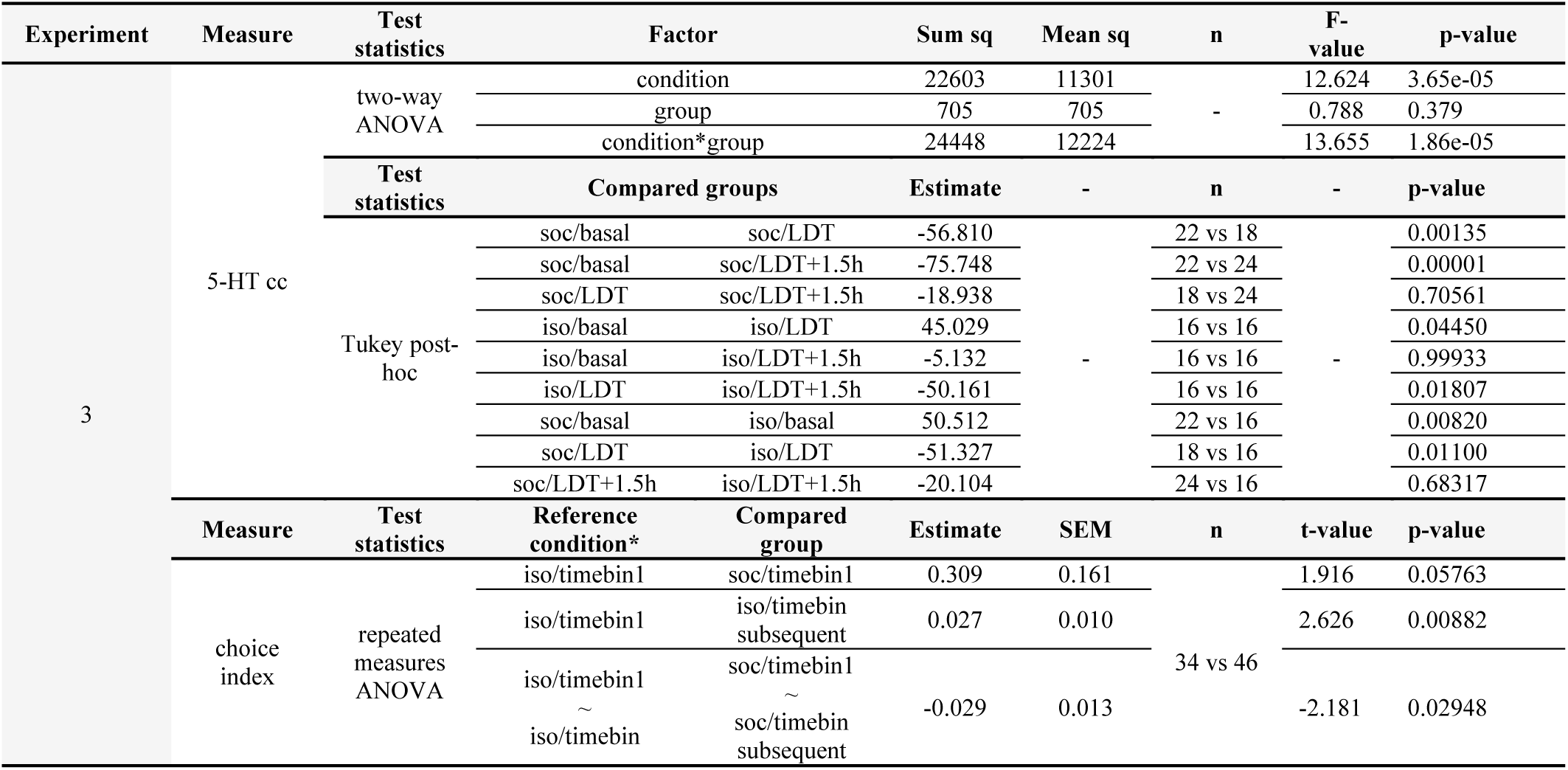
Statistical data of *Experiment 3*. Note that different hypotheses testing methods were used, hence shown parameters may differ from method to method.

**Table 4:**
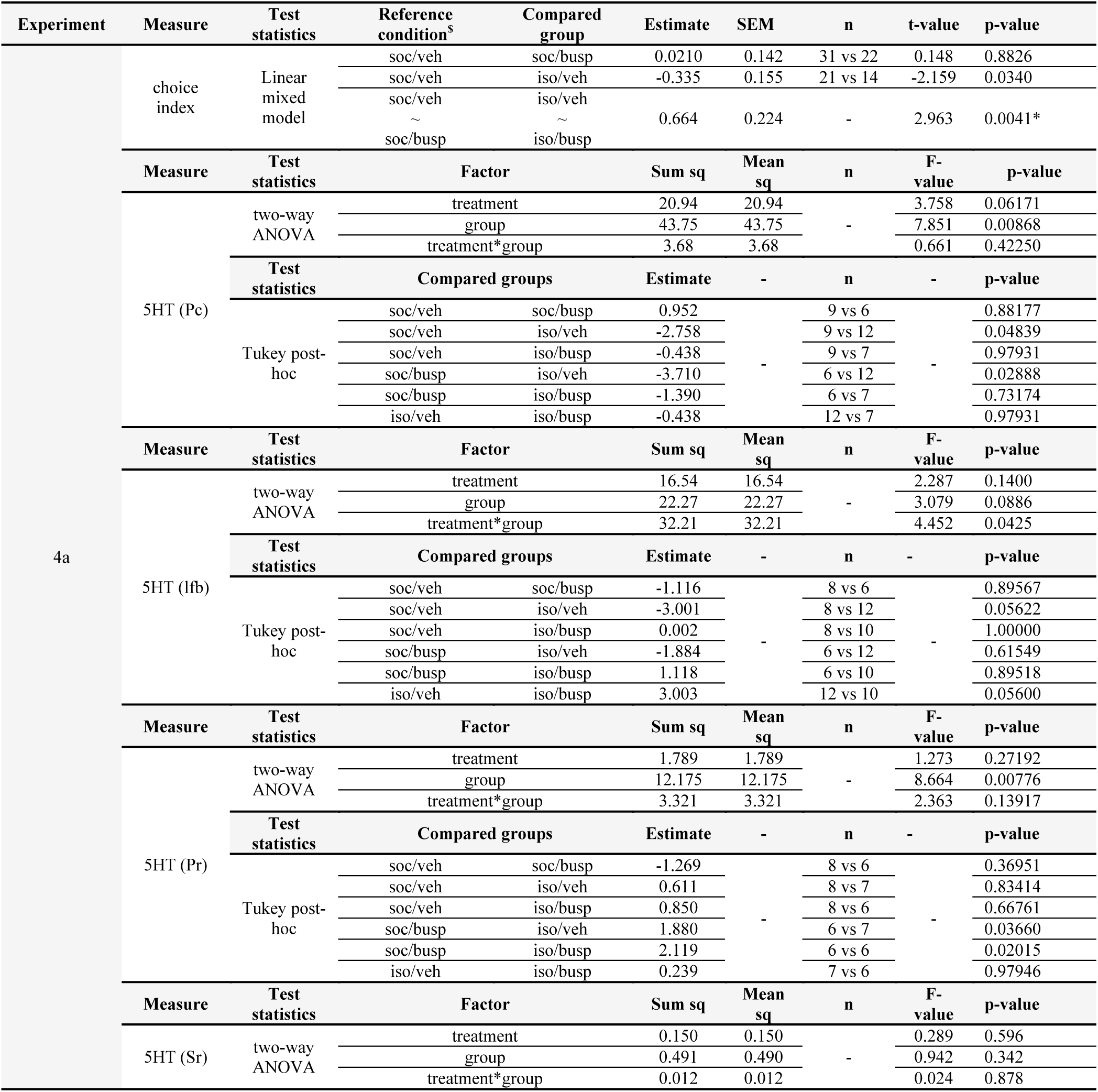
Statistical data of *Experiment 4a*. ^$^ The statistical output of these linear mixed models are able to show significant differences from a reference condition in each analyzed levels, i.e. rearing and treatment. Rearing “social” and treatment “vehicle” were set as reference conditions. We also show interactional effects in relation to the reference categories, e.g. the difference between the treatment vehicle and buspirone (∼) in socially reared groups differ from the corresponding difference between the isolated groups. Note that different hypotheses testing methods were used, hence shown parameters may differ from method to method.

**Table 5:**
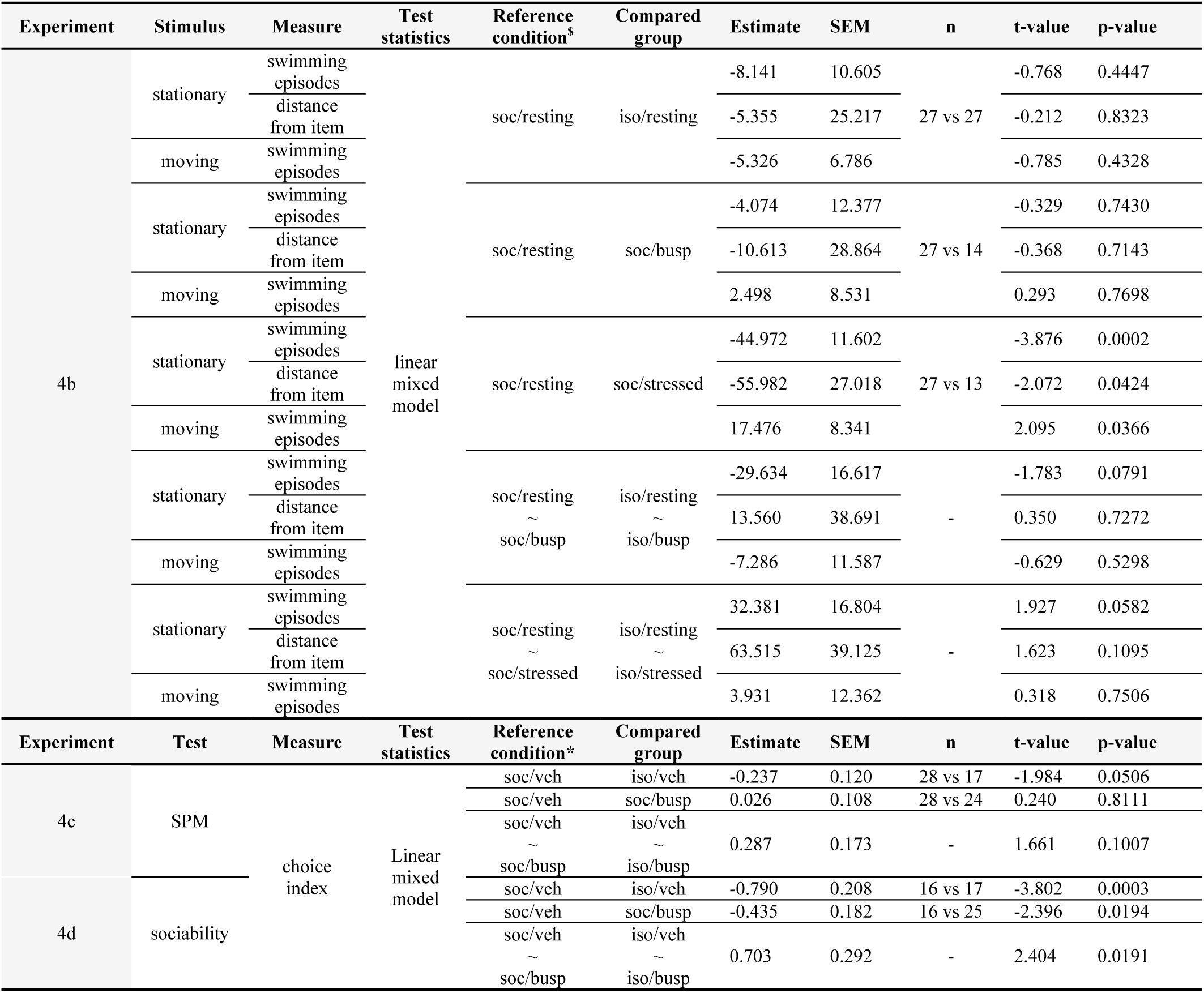
Statistical data of *Experiment 4b*, *4c* and *4d*. ^$^ The statistical output of these linear mixed models are able to show significant differences from a reference condition in each analyzed levels, i.e. rearing and treatment/condition. Rearing “social” and treatment/condition “resting” were set as reference conditions. We also show interactional effects in relation to the reference categories, e.g. the difference between the resting and novelty condition (∼) in socially reared groups do not differ from the corresponding difference between the isolated groups. Note that different hypotheses testing methods were used, hence shown parameters may differ from method to method.

### Social isolation-induced 5-HT changes are specific to the dorsal pallium and, along with behavioral deficits, can be prevented by blockade of 5-HT signaling

In *Experiment 4a*, we assessed the functional connectivity between isolation-induced behavioral and physiological effects. We administered buspirone prior to the post-metamorphic LDT test and used immunohistochemical labeling for 5-HT in several regions of the telencephalon as well (Figure 4c). We hypothesized that such a pharmacological treatment might possibly be able to dampen 5-HT signaling in forebrain areas relevant to the regulation of behavioral processes, since i) buspirone predominantly acts on 5HT1A receptors (Loane, 2012), ii) zebrafish express these mainly as autoreceptors of the raphe nuclei (Norton et al., 2008), which are iii) the only serotonergic cell clusters that project to the teleostean telencephalon (Maximino et al., 2013a), iv) the area that orchestrates dark avoidance behavior (Lau and Guo, 2011). Furthermore, v) buspirone binds to 5HT1A receptors of zebrafish with a 4 times greater affinity than that of rats and an 8 times greater affinity than that of humans (Barba-Escobedo and Gould, 2012). Isolated fish, similarly to those in *Experiment 2* and *3*, showed decreased light avoidance behavior. This effect was preventable by an acute buspirone treatment with a dose that had no effect on socially reared conspecifics (Figure 4b). Immunohistochemical labeling revealed higher 5-HT immunoreactivity in isolated fish compared to controls in the lateral forebrain bundle and in the caudal, but not in any rostral part of the pallium. This difference was abolished by the acute administration of buspirone (Figure 4d and 4e). Taken together, our results display that with a single 5HT1AR agonist treatment we were able to restore the normal novelty-induced avoidance phenotype. In addition, we found 5-HT level differences in fibers that enter the forebrain and target the caudal recess of the telencephalon where cell masses of the dorsomedial and dorsolateral areas, homologue structures of the mammalian amygdala and hippocampus, emerge separately (Wulliman et al., 2012). Such differences were preventable by acute buspirone treatment. In summary, we detected functional connections between forebrain-localized 5-HT response and inadequate avoidance behavior.

### Social isolation shifts response capacity between novel and resting conditions and is inverted by the effects of 5-HT signaling blockade

Due to the highly different levels of 5-HT in resting and novel environment, including changes in limbic structures, and ineffective challenge responding of isolated fish, in *Experiment 4b* we assessed possible isolation-induced impairment in stress-induced arousal state. Alertness in mammals is associated with activation of the amygdala (Gläscher and Adolphs, 2003; Harmer et al., 2006) and modulated by 5-HT signaling (Spoont, 1992), hence we speculated that an inadequate emergence of such condition may drive the isolation-induced behavioral impairment of zebrafish. In order to assess this issue, a stationary black slider was projected on the bottom of testing compartments for 5 minutes, which subsequently started to move in circles with increasing speed (Figure 5a). We sampled behavioral responses in resting conditions, after 20 minutes of habituation, as well as in novel environment, immediately after the transfer to the apparatus. Normally, an animal in novelty-induced alert state react more rapidly to movement or react to a yet slower moving objects (Yokogawa et al., 2012) compared to habituated resting individuals. Socially reared animals, in novel but not in familiar context, showed a decrease in the number of swimming episodes and in the distance from the stationary slider in response to its appearance, while buspirone prevented both of these effects (Figure 5b). Interestingly, isolated fish only showed such stimuli-induced decreased locomotion in response to buspirone treatment. In response to the movement stimulus, socially reared animals increased the frequency of their swimming episodes in novel, but not in resting conditions, while buspirone abolished this reaction (Figure 5c). In contrast, isolated fish showed higher response frequency in resting, and a delayed, oppositely directed response in stress-induced conditions. Buspirone pre-treatment of isolated fish triggered a response similar in shape that of socially reared stressed or isolated habituated animals. In summary, we detected an isolation-induced lack of typical stress-induced, arousal-associated activity in novel, and an exacerbated activity in resting conditions, while buspirone evoked opposite direction effects compared to socially reared animals.

**Figure 5:**
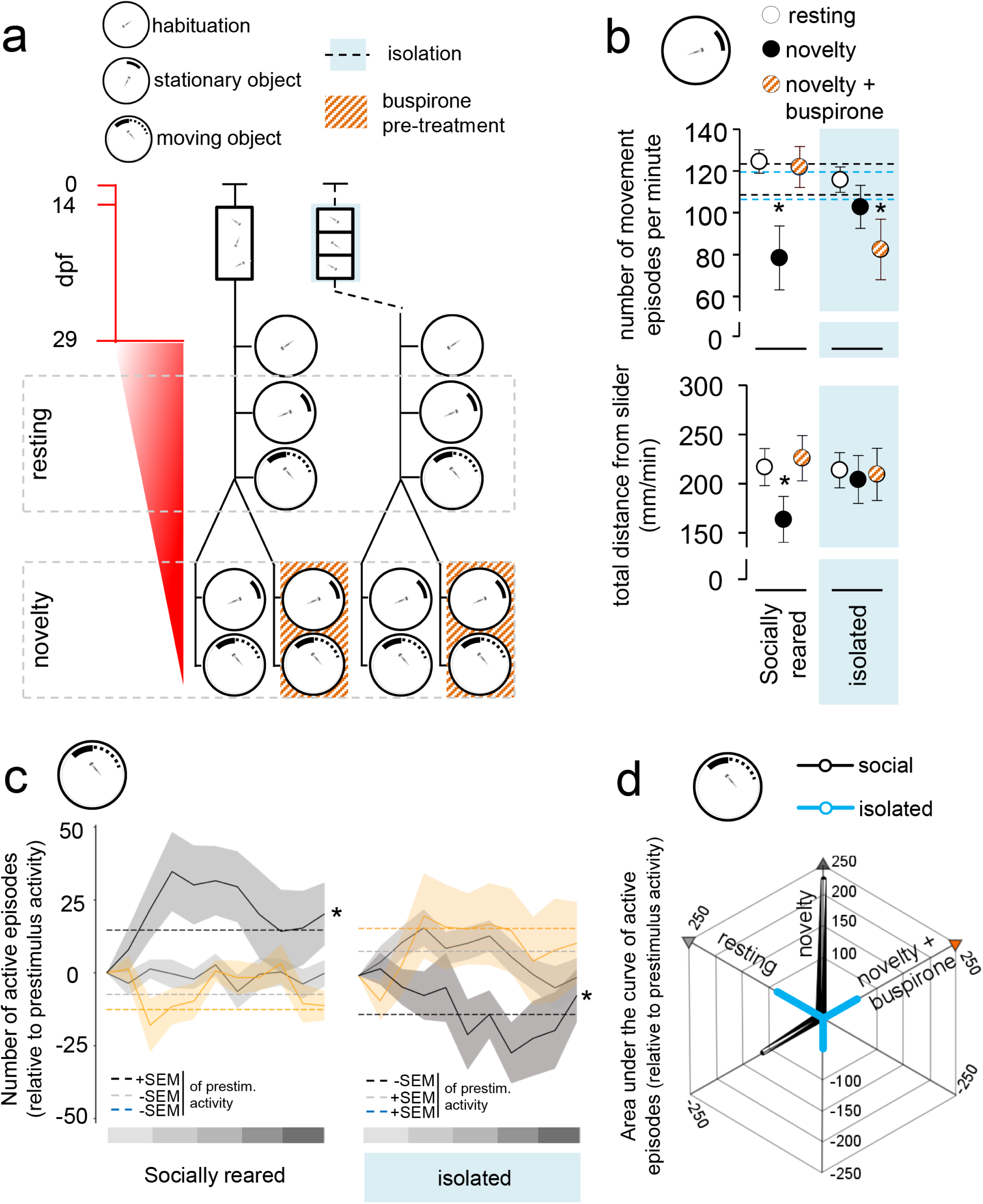
**a)** Protocol of *Experiment 4b*. Red timeline shows the age of larvae expressed in days post fertilisation (dpf). **b)** Locomotor response to (top) and the total distance from (bottom) the projected stationary object in resting and novel conditions and in response to prior buspirone treatment. Reference levels of the linear mixed model were „social rearing” and „condition resting”. Asterisk indicates differences between experimental conditions (resting, novelty, novelty + buspirone) at the level of each rearing conditons. Dasehed lines indicate ± standard error of mean of activity during the habituation period. **c)** Minute-by-minute resolution analysis of locomotor response to the moving object in resting and novel conditons and in response to prior buspirone treatment. The number of active swimming episodes are presented relative to previous (prestim) activity of the individuals, band shows ± standard error of mean. Grey gradient scale represents the dynamics of speed increase of the moving object. Dashed lines mark standard errors of means of the prestimulus activity. Asterisk represents significant differences from resting condition. **d)** Arousal profiles of socially reared and isolated animals presented as locomotor responses to the moving object under different conditions (resting, novelty, novelty + buspirone). 3 dimension starplot shows the 3 different conditions. Locomotor response is presented as the area under the curve of active episodes relative to the previous activity of the individuals.

### Social isolation-induced decrease in responsiveness is specific to novelty-based visually-driven challenges

Given such isolation-induced low responsivity, potentially driven by an inability to form stress-associated arousal, next we investigated whether the effects are challenge-specific. In order to assess this issue, in Experiment 4c and 4d, we analyzed isolation-induced changes in response to different environmental challenges, such as the swimming plus-maze (Varga et al., 2018) and the U-shaped social preference tests (Dreosti et al., 2015). Similarly to the LDT test, both tests trigger approach-avoidance conflict in the animal, however, presenting different type of stimuli, e.g. non-visual non-meaningful (novelty) and visual meaningful (social) challenges. To analyze the potential role of 5-HT signaling as well, we administered buspirone prior to the tests (Figure 6a). Interestingly, isolated fish showed increased avoidance compared to socially reared controls in both tests, however, buspirone completely (Figure 6b) or partially (Figure 6c) prevented such effects. Note, that in the case of the social preference test, buspirone abolished social activity regardless of its direction, e.g. the preference in case of socially reared and the aversion in case of isolated animals. These results indicate that decreased responsiveness of isolated animals is specific to visual non-meaningful, novel stimuli, e.g. the light/dark or the sensory-motor reaction tests.

**Figure 6:**
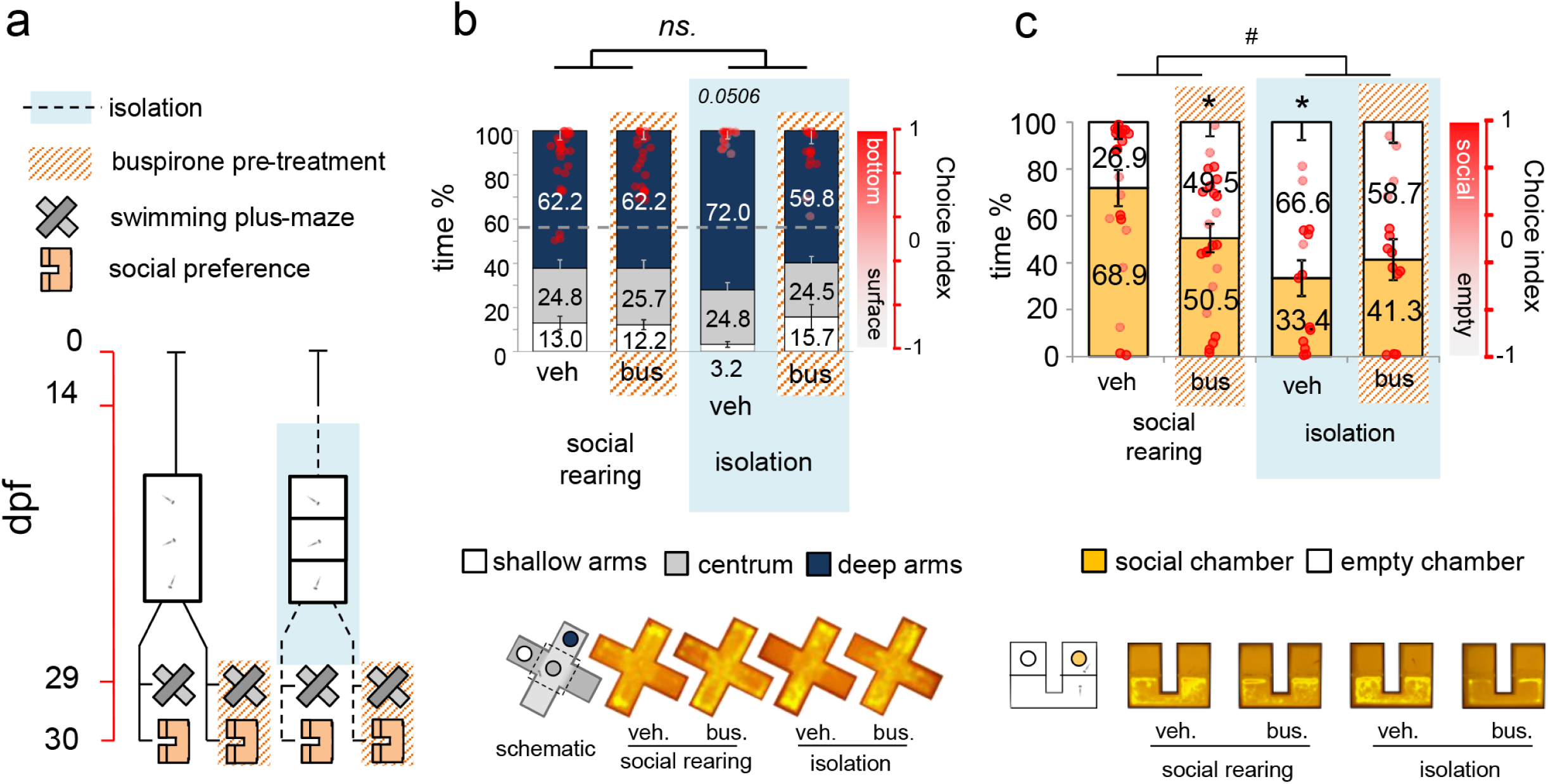
**a)** Protocol of *Experiment 4c* and *4d*. Red timeline shows the age of larvae expressed in days post fertilisation (dpf). **b)** Surface avoidance behavior of post-metamorphic zebrafish 2 weeks following chronic social isolation and/or acute buspirone treatment (top) and their representative heatmaps (bottom). Left axis shows the percentage of time, right axis shows choice index in which 1 indicates 100% time in the deep zone, whereas −1 represents 100% time in the shallow zone. The reference levels of the linear mixed model were „social rearing” and „treatement vehicle”. **c)** Social preference of post-metamorphic zebrafish 2 weeks after chronic social isolation and/or after buspirone treatment (top) and their representative heatmaps (bottom). Left axis shows the percentage of time, right axis shows choice index in which 1 indicates 100% time in the social chamber, whereas −1 represents 100% time in the empty chamber. The reference levels of the linear mixed model were „social rearing” and „treatement vehicle”. Asterisk indicates significant differences from socially reared vehicle treated animals, hashtag indicates significant differences between the relation of the treatment levels in socially reared and isolated groups.

## Discussion

We have characterized a period during zebrafish development that is highly reactive to the environment in terms of the ability of the individual to change behavioral responses to a repeated challenge. Absence of social stimuli during this period promoted an atypical challenge-coping phenotype, characterized by exacerbated alertness under resting conditions and inability to show novelty-induced arousal and avoidance behavior. In line with these behavioral impairments, isolated fish expressed lower baseline, and same magnitude, but oppositely directed serotonin-level changes in response to novelty compared to socially reared subjects. This effect was localized in the caudal excess of the dorsal pallium, a brain area responsible for adequate stress-induced aversion. 5HT1AR agonist treatment prevented isolation-induced serotonin response in the forebrain and rescued physiological challenge-induced behaviors as well, indicating a functional connection between these processes. The effects were specific to visual challenges in a novel context, since they were undetectable at non-visual novelty, and visual social challenges.

In our first experiments, we showed that a switch from dark to light avoidance, a specific aspect of the behavioral metamorphosis, occurs between the first and second week of development in zebrafish. Furthermore, we showed that this process overlaps with the ability to decrease avoidance in response to repeated challenge, suggesting that this switch is accompanied by enhanced behavioral plasticity, thus it may serve as a tool to model early life experience-dependent phenomena. The light/dark choice reversal was first observed by Lau and Guo (2011), who estimated its occurrence later than we did, between the second and third week of development. This discrepancy indicates a potential experience-dependent flexibility in the onset of such process, while such variability in the onset supports our hypothesis that the first month of ontogenesis possibly represents a highly plastic, reactive developmental phase. Decreased avoidance at repeated testing supports this idea as well, as it may be driven by habituation, a neuroplasticity-based process (Thompson and Spencer, 1966). The idea of habituation is supported by our findings, as i) decreased avoidance was detectable regardless of the direction of baseline preference, ii) it was not caused by aging, and iii) stronger baseline preference predicted a stronger change in its expression. In summary, one aspect of the behavioral metamorphosis coincides with and speculatively requires a peak in adaptive capacity.

In our next experiment, we showed that an acute challenge, in contrast to its short-term effect, does not influence avoidance 2 weeks later, indicating a spontaneous recovery from novelty stress during the long inter-test period. In contrast, absence of social stimuli during this sensitive period exerted robust effects on behavior, including decreased avoidance and locomotion. Since the LDT test, used here to mimic an acute environmental challenge, is a pharmacologically (Chen et al., 2015; Gould, 2011; Maximino, 2011), physiologically (Chen et al., 2015) and behaviorally (Blaser, 2011) validated tool for measuring anxiety-associated behavior of zebrafish, such decreased avoidance might be interpreted as social isolation-induced anxiolysis. This theory, however, is unlikely, as i.) application of the anxiolytic agent buspirone prevented the decrease, thus, if the ‘anxiolysis hypothesis’ is accepted, it acted as an anxiogenic drug, contradicting its widely accepted effect on behavior; ii.) phenomenological correlates of other anxiety-like domains were enhanced in response to social isolation. According to rodent and zebrafish literature, it is rare, but not unique (Parker, 2012; Shams et al., 2015), that such a chronic environmental perturbation as social isolation promotes a low anxiety phenotype (for reviews see Fone and Porkess, 2008; Hall, 1998; Lukkes et al., 2009), particularly, several authors discussed social isolation of zebrafish as beneficial, since it consistently decreased stress-induced cortisol response and in some cases avoidance as well (Forsatkar et al., 2017; Giacomini, 2015; Parker, 2012; Shams et al., 2015). Despite the importance of these results, we suggest a more careful interpretation of the above data. Clinical and preclinical data suggest that an inadequate cortisol response may drive or at least coincide with abnormal behavior, regardless of the direction in which it moves out of the physiological range (Gutiérrez et al., 2019; Haller et al., 2001; McBurnett et al., 2000; Toth et al., 2011; van Goozen et al., 1998; Vanyukov et al., 1993). Furthermore, the corticosteroid response is not necessarily in a positive or in any relation with the expression of anxiety-like behavior (Aliczki et al., 2013; Mikics et al., 2005). On the other hand, isolation-induced low avoidance has also been interpreted as reduced pro-active coping and described as a stable “slow-emerging” phenotype in rodents (Arakawa, 2007, 2005). Isolated rats consistently showed a higher latency to enter the centrum of an unfamiliar open field arena, or a lower frequency of entries into the centrum of a familiar but brightly lit open field arena, which indicates that isolation-induced delayed exploration is related to adversity and the accompanying stress-response. In another study, socially reared zebrafish individuals showing delayed emergence from a dark to a brightly lit arena, a phenotype resembling the one observed in our study, showed lower locomotion in general and lower challenge-induced but similar baseline cortisol responses (Tudorache et al., 2015). One can hypothesize that a low stress-induced cortisol response, without any change in basal levels, and the accompanying low avoidance behavior might indicate a malfunction of challenge-responding. This phenotypic background may account for low avoidance in adult and decreased locomotion in 30 dpf juvenile zebrafish after social isolation as well, observed by Shams and colleagues (Shams et al., 2017a, 2015). This theory is also supported by our findings on isolation-induced low, delayed avoidance and inadequate arousal in response to a challenge and enhanced arousal under baseline, resting conditions. This indicates that isolated fish shift stress-associated alertness to baseline conditions and in this manner are unable to develop adequate internal states and accompanying behavioral responsiveness in neither of the contexts. This also means that lack of stimuli during the characterized, highly plastic developmental period has a crucial role in shaping the post-metamorphic behavioral phenotype.

Given the low responding behavioral phenotype, we assessed isolation-induced effects on 5-HT signaling since it is highly sensitive to early life adversity and is known as a prominent modulator of mood, as well as general or specific arousal states (Lucki, 1998; Soubrié, 1986; Spoont, 1992). We found that isolated fish show decreased baseline 5-HT tone and, contrarily to socially reared conspecifics, a rapid gradual 5-HT increase in response to the LDT challenge. This difference was non-significant 1.5 hours later on a whole-brain level, however, it was still detectable in fibers projecting to pallial areas, e.g. the lateral forebrain bundle and the caudal recess of the dorsal pallium. Furthermore, 5HT1AR agonist treatment prevented differences in forebrain 5-HT immunoreactivity and behavior as well. Our findings regarding brain 5-HT levels are in contradiction with previous reports, however, this discrepancy might be explained by the facts that in these papers either i.) only extracellular 5-HT content was measured and ii) adult animals were employed (Maximino et al., 2013b) or iii) social isolation was conducted in adult fish or in larvae from the age of 0 dpf (Shams et al., 2017a, 2017b, 2015). The latter case, together with rodent data supports that the period during which isolation occurs is fundamental regarding later consequences, highlighting the importance of critical and sensitive developmental periods (Einon and Morgan, 1977; Fone and Porkess, 2008; Hall, 1998; Lukkes et al., 2009a). We suggest that the environmental perturbation we applied in a period characterized by enhanced adaptive capacity exerts neurochemical and behavioral effects that overtop the ones previously observed by other authors. It is also important to note that besides the discrepancy between our results and data from other laboratories employing zebrafish models, numerous clinical and preclinical rodent studies support our findings: several authors described 5-HT tone as a general inhibitor of responsiveness, which decreases in response to sudden environmental changes allowing the expression of adequate behavioral responses. From this perspective, 5-HT acts as a context-dependent modulator that levels responsiveness into non-impulsive ranges (Brodie and Shore, 1957; Depue and Spoont, 1986; Lucki, 1998; Soubrié, 1986; Spoont, 1992). Behrendt suggests that 5-HT downregulates responsivity partially through its actions on amygdala function (Ralf-Peter Behrendt, 2011), which theory is supported by several human fMRI studies investigating the mechanisms of selective 5-HT reuptake inhibitors in aversive contexts (Del-Ben et al., 2005; Harmer et al., 2006; Murphy et al., 2009). In addition, isolation-induced enhanced serotonergic activity and fiber density in the amygdala were also reported in most rodent neurochemistry studies (Gos et al., 2006; Günther et al., 2008; Lehmann et al., 2003), in line with our current results.

In summary, we suggest that 5-HT tone of zebrafish restricts behavior under resting conditions, while its phasic decrease yields behavioral stress responses during a challenge. Social isolation during an adaptive time-window leads to an abnormal 5-HT phenotype in juveniles, with serotonin levels inversely reflecting those of socially reared ones, thus exerting shifted responsivity as well. The phenomenon is modulated in the caudal recess of the telencephalon, where homologue structures of the mammalian amygdala and hippocampus are located. Our findings indicate that zebrafish possess a conserved serotonergic mechanism that context-dependently modulates environmental reactivity and is highly sensitive to early life experiences.

## Acknowledgement

We thank Christina Miskolczi (M.Sc., IEM, HAS, Budapest, Hungary) for language editing and Máté Tóth (Ph.D., IEM, HAS, Budapest, Hungary) for sharing comments that greatly improved the study. We also thank László Barna and Csaba Pongor, the Nikon Microscopy Center at the Institute of Experimental Medicine, Nikon Austria GmbH, and Auro-Science Consulting, Ltd., for kindly providing microscopy support.

## References

Aliczki, M., Zelena, D., Mikics, E., Varga, Z.K., Pinter, O., Bakos, N.V., Varga, J., Haller, J., 2013. Monoacylglycerol lipase inhibition-induced changes in plasma corticosterone levels, anxiety and locomotor activity in male CD1 mice. Horm. Behav. 63, 752–758.

András Péter, 2017. Solomon Coder.

Arakawa, H., 2007. Ontogeny of sex differences in defensive burying behavior in rats: effect of social isolation. Aggress. Behav. 33, 38–47.

Arakawa, H., 2005. Interaction between isolation rearing and social development on exploratory behavior in male rats. Behav. Processes 70, 223–234.

Barba-Escobedo, P.A., Gould, G.G., 2012. Visual social preferences of lone zebrafish in a novel environment: strain and anxiolytic effects. Genes Brain Behav. 11, 366–373.

Barret Schloerke, Jason Crowley, Di Cook, Francois Briatte, Moritz Marbach, Edwin Thoen, Amos Elberg, Joseph Larmarange, 2017. GGally: Extension to “ggplot2.”

Blaser, R.E., 2011. Stimuli affecting zebrafish (Danio rerio) behavior in the light/dark preference test. Physiol. Behav. 831–837.

Booij, L., Tremblay, R.E., Szyf, M., Benkelfat, C., 2015. Genetic and early environmental influences on the serotonin system: consequences for brain development and risk for psychopathology. J. Psychiatry Neurosci. JPN 40, 5–18.

Brodie, B.B., Shore, P.A., 1957. A CONCEPT FOR A ROLE OF SEROTONIN AND NOREPINEPHRINE AS CHEMICAL MEDIATORS IN THE BRAIN. Ann. N. Y. Acad. Sci. 66, 631–642.

Chapman, D.P., Whitfield, C.L., Felitti, V.J., Dube, S.R., Edwards, V.J., Anda, R.F., 2004. Adverse childhood experiences and the risk of depressive disorders in adulthood. J. Affect. Disord. 82, 217–225.

Chen, F., Chen, S., Liu, S., Zhang, C., Peng, G., 2015. Effects of lorazepam and WAY-200070 in larval zebrafish light/dark choice test. Neuropharmacology 95, 226–233.

Del-Ben, C.M., Deakin, J.F.W., McKie, S., Delvai, N.A., Williams, S.R., Elliott, R., Dolan, M., Anderson, I.M., 2005. The effect of citalopram pretreatment on neuronal responses to neuropsychological tasks in normal volunteers: an FMRI study. Neuropsychopharmacol. Off. Publ. Am. Coll. Neuropsychopharmacol. 30, 1724–1734.

Depue, R.A., Spoont, M.R., 1986. A Behavioral Dimension of Constraint. Ann. N. Y. Acad. Sci. 487, 47–62. https://doi.org/10.1111/j.1749-6632.1986.tb27885.x

Douglas Bates, 2015. Fitting Linear Mixed-Effects Models Using {lme4}. J. Stat. Softw. 67, 1–48.

Dreosti, E., Lopes, G., Kampff, A.R., Wilson, S.W., 2015. Development of social behavior in young zebrafish. Front. Neural Circuits 9.

Egan, R.J., Kalueff, A.V., 2009. Understanding behavioral and physiological phenotypes of stress and anxiety in zebrafish. Behav. Brain Res. 38–44.

Einon, D.F., Morgan, M.J., 1977. A critical period for social isolation in the rat. Dev. Psychobiol. 10, 123–132.

Fone, K.C.F., Porkess, M.V., 2008. Behavioural and neurochemical effects of post-weaning social isolation in rodents—Relevance to developmental neuropsychiatric disorders. Neurosci. Biobehav. Rev., The long-term consequences of stress on brain function: from adaptation to mental diseases 32, 1087–1102.

Forsatkar, M.N., Safari, O., Boiti, C., 2017. Effects of social isolation on growth, stress response, and immunity of zebrafish. Acta Ethologica 20, 255–261.

Geyer, M.A., 1995. Serotonergic functions in arousal and motor activity. Behav. Brain Res., Serotonin 73, 31–35.

Giacomini, A.C.V.V., 2015. My stress, our stress: Blunted cortisol response to stress in isolated housed zebrafish. Physiol. Behav. 182–187.

Gläscher, J., Adolphs, R., 2003. Processing of the Arousal of Subliminal and Supraliminal Emotional Stimuli by the Human Amygdala. J. Neurosci. 23, 10274–10282.

Gos, T., Becker, K., Bock, J., Malecki, U., Bogerts, B., Poeggel, G., Braun, K., 2006. Early neonatal and postweaning social emotional deprivation interferes with the maturation of serotonergic and tyrosine hydroxylase-immunoreactive afferent fiber systems in the rodent nucleus accumbens, hippocampus and amygdala. Neuroscience 140, 811–821.

Gould, GeorgiannaG., 2011. Aquatic Light/Dark Plus Maze Novel Environment for Assessing Anxious Versus Exploratory Behavior in Zebrafish (Danio rerio) and Other Small Teleost Fish, in: Kalueff, A.V., Cachat, J.M. (Eds.), Zebrafish Neurobehavioral Protocols, Neuromethods. Humana Press, pp. 99–108.

Gross, C., Zhuang, X., Stark, K., Ramboz, S., Oosting, R., Kirby, L., Santarelli, L., Beck, S., Hen, R., 2002. Serotonin1A receptor acts during development to establish normal anxiety-like behaviour in the adult. Nature 416, 396–400.

Günther, L., Liebscher, S., Jähkel, M., Oehler, J., 2008. Effects of chronic citalopram treatment on 5-HT1A and 5-HT2A receptors in group- and isolation-housed mice. Eur. J. Pharmacol. 593, 49–61.

Gutiérrez, H.C., Colanesi, S., Cooper, B., Reichmann, F., Young, A.M.J., Kelsh, R.N., Norton, W.H.J., 2019. Endothelin neurotransmitter signalling controls zebrafish social behaviour. Sci. Rep. 9, 3040.

Hall, F.S., 1998. Social Deprivation of Neonatal, Adolescent, and Adult Rats Has Distinct Neurochemical and Behavioral Consequences. Crit. Rev. Neurobiol. 12.

Haller, J., Harold, G., Sandi, C., Neumann, I.D., 2014. Effects of Adverse Early-Life Events on Aggression and Anti-Social Behaviours in Animals and Humans. J. Neuroendocrinol. 26, 724–738.

Haller, J., Schraaf, J.V.D., Kruk, M.R., 2001. Deviant Forms of Aggression in Glucocorticoid Hyporeactive Rats: A Model for ‘Pathological’ Aggression? J. Neuroendocrinol. 13, 102–107.

Harmer, C.J., Mackay, C.E., Reid, C.B., Cowen, P.J., Goodwin, G.M., 2006. Antidepressant drug treatment modifies the neural processing of nonconscious threat cues. Biol. Psychiatry 59, 816–820.

Jing, J., Gillette, R., Weiss, K.R., 2009. Evolving concepts of arousal: insights from simple model systems. Rev. Neurosci. 20, 405–427.

Julian J. Faraway, 2016. Extending the Linear Model with R: Generalized Linear, Mixed Effects and Nonparametric Regression Models. CRC Press.

Kuznetsova, A., Per Bruun Brockhoff, Rune Haubo Bojesen Christensen, 2016. lmerTest: Tests in Linear Mixed Effects Models.

Lau, B.Y.B., Guo, S., 2011. Identification of a brain center whose activity discriminates a choice behavior in zebrafish. PNAS 108, 2581–2586.

Lehmann, K., Lesting, J., Polascheck, D., Teuchert-Noodt, G., 2003. Serotonin fibre densities in subcortical areas: differential effects of isolated rearing and methamphetamine. Brain Res. Dev. Brain Res. 147, 143–152.

Lieschke, P.D., Currie, G.J., 2007. Animal models of human disease: zebrafish swim into view. Nat. Rev. Genet.

Loane, C., 2012. Buspirone: What is it all about? Brain Res. 111–118.

Lovett-Barron, M., Andalman, A.S., Allen, W.E., Vesuna, S., Kauvar, I., Burns, V.M., Deisseroth, K., 2017. Ancestral Circuits for the Coordinated Modulation of Brain State. Cell 171, 1411–1423.e17.

Lucki, I., 1998. The spectrum of behaviors influenced by serotonin. Biol. Psychiatry 44, 151– 162.

Lukkes, J.L., Summers, C.H., Scholl, J.L., Renner, K.J., Forster, G.L., 2009a. Early life social isolation alters corticotropin-releasing factor responses in adult rats. Neuroscience 158, 845– 855.

Lukkes, J.L., Watt, M.J., Lowry, C.A., Forster, G.L., 2009b. Consequences of Post-Weaning Social Isolation on Anxiety Behavior and Related Neural Circuits in Rodents. Front. Behav. Neurosci. 3.

MacPhail, R.C., 2009. Locomotion in larval zebrafish: Influence of time of day, lighting and ethanol. Neurotoxicology 52–58.

MacRae, C.A., Peterson, R.T., 2015. Zebrafish as tools for drug discovery. Nat. Rev. Drug Discov. 14, 721–731.

Maximino, C., 2011. Pharmacological analysis of zebrafish (Danio rerio) scototaxis. Neuropsychopharmacol. Biol. Psychiatry 624–631.

Maximino, C., Lima, M.G., Araujo, J., Oliveira, K.R.M., Herculano, A.M., Stewart, A.M., Kyzar, E.J., Cachat, J., Kalueff, A.V., 2013a. The serotonergic system of zebrafish: genomics, neuroanatomy and neuropharmacology. Serotonin Biosynth. Regul. Health Implic. Nova Sci. N. Y. 53–67.

Maximino, C., Puty, B., Benzecry, R., Araújo, J., Lima, M.G., de Jesus Oliveira Batista, E., Renata de Matos Oliveira, K., Crespo-Lopez, M.E., Herculano, A.M., 2013b. Role of serotonin in zebrafish (Danio rerio) anxiety: Relationship with serotonin levels and effect of buspirone, WAY 100635, SB 224289, fluoxetine and para-chlorophenylalanine (pCPA) in two behavioral models. Neuropharmacology 71, 83–97.

McBurnett, K., Lahey, B.B., Rathouz, P.J., Loeber, R., 2000. Low salivary cortisol and persistent aggression in boys referred for disruptive behavior. Arch. Gen. Psychiatry 57, 38– 43.

Mikics, E., Barsy, B., Barsvári, B., Haller, J., 2005. Behavioral specificity of non-genomic glucocorticoid effects in rats: effects on risk assessment in the elevated plus-maze and the open-field. Horm. Behav. 48, 152–162.

Murphy, S.E., Norbury, R., O’Sullivan, U., Cowen, P.J., Harmer, C.J., 2009. Effect of a single dose of citalopram on amygdala response to emotional faces. Br. J. Psychiatry J. Ment. Sci. 194, 535–540.

Nederhof, E., Schmidt, M.V., 2012. Mismatch or cumulative stress: Toward an integrated hypothesis of programming effects. Physiol. Behav., Special Section: The Mismatch Hypothesis of Psychiatric Disease 106, 691–700.

Noldus L.P.J.J., Spink A.J., Tegelenbosch R.A.J., 2001. EthoVision: A versatile video tracking system for automation of behavioral experiments. Behav. Res. Methods Instrum. Comput. 3, 398–414.

Norton, W.H.J., Folchert, A., Bally-Cuif, L., 2008. Comparative analysis of serotonin receptor (HTR1A/HTR1B families) and transporter (*slc6a4a/b*) gene expression in the zebrafish brain. J. Comp. Neurol. 511, 521–542.

Panula, P., Chen, Y.-C., Priyadarshini, M., Kudo, H., Semenova, S., Sundvik, M., Sallinen, V., 2010. The comparative neuroanatomy and neurochemistry of zebrafish CNS systems of relevance to human neuropsychiatric diseases. Neurobiol. Dis., Special issue: Non-mammalian models of neuropsychiatric disease 40, 46–57.

Parker, M.O., 2012. Housing Conditions Differentially Affect Physiological and Behavioural Stress Responses of Zebrafish, as well as the Response to Anxiolytics. Plos One 7.

R Core Team, 2017. R: A Language and Environment for Statistical Computing. R Foundation for Statistical Computing, Vienna, Austria.

Raleigh, M.J., McGuire, M.T., Brammer, G.L., Yuwiler, A., 1984. Social and environmental influences on blood serotonin concentrations in monkeys. Arch. Gen. Psychiatry 41, 405–410.

Ralf-Peter Behrendt, 2011. Neuroanatomy of Social Behaviour_ An Evolutionary and Psychoanalytic Perspective.

Santarelli, S., Lesuis, S.L., Wang, X.-D., Wagner, K.V., Hartmann, J., Labermaier, C., Scharf, S.H., Müller, M.B., Holsboer, F., Schmidt, M.V., 2014. Evidence supporting the match/mismatch hypothesis of psychiatric disorders. Eur. Neuropsychopharmacol. 24, 907– 918.

Schneider, C.A., Rasband, W.S., Eliceiri, K.W., 2012. NIH Image to ImageJ: 25 years of image analysis. Nat. Methods 9, 671–675.

Shams, S., Amlani, S., Buske, C., Chatterjee, D., Gerlai, R., 2017a. Developmental social isolation affects adult behavior, social interaction, and dopamine metabolite levels in zebrafish. Dev. Psychobiol.

Shams, S., Chatterjee, D., Gerlai, R., 2015. Chronic social isolation affects thigmotaxis and whole-brain serotonin levels in adult zebrafish. Behav. Brain Res. 292, 283–287.

Shams, S., Seguin, D., Facciol, A., Chatterjee, D., Gerlai, R., 2017b. Effect of social isolation on anxiety-related behaviors, cortisol, and monoamines in adult zebrafish. Behav. Neurosci. 131, 492–504.

Soubrié, P., 1986. Reconciling the role of central serotonin neurons in human and animal behavior [WWW Document]. Behav. Brain Sci.

Spoont, M.R., 1992. Modulatory role of serotonin in neural information processing: Implications for human psychopathology. Psychol. Bull. 112, 330–350.

Thompson, R.F., Spencer, W.A., 1966. Habituation: a model phenomenon for the study of neuronal substrates of behavior. Psychol. Rev. 73, 16–43.

Toth, M., Mikics, E., Tulogdi, A., Aliczki, M., Haller, J., 2011. Post-weaning social isolation induces abnormal forms of aggression in conjunction with increased glucocorticoid and autonomic stress responses. Horm. Behav. 60, 28–36.

Tudorache, C., ter Braake, A., Tromp, M., Slabbekoorn, H., Schaaf, M.J.M., 2015. Behavioral and physiological indicators of stress coping styles in larval zebrafish. Stress Amst. Neth. 18, 121–128.

van Goozen, S.H., Matthys, W., Cohen-Kettenis, P.T., Gispen-de Wied, C., Wiegant, V.M., van Engeland, H., 1998. Salivary cortisol and cardiovascular activity during stress in oppositional-defiant disorder boys and normal controls. Biol. Psychiatry 43, 531–539.

Vanyukov, M.M., Moss, H.B., Plail, J.A., Blackson, T., Mezzich, A.C., Tarter, R.E., 1993. Antisocial symptoms in preadolescent boys and in their parents: associations with cortisol. Psychiatry Res. 46, 9–17.

Varga, Z.K., Zsigmond, Á., Pejtsik, D., Varga, M., Demeter, K., Mikics, É., Haller, J., Aliczki, M., 2018. The swimming plus-maze test: a novel high-throughput model for assessment of anxiety-related behaviour in larval and juvenile zebrafish (Danio rerio). Sci. Rep. 8, 16590.

Westerfield, M., 2000. The Zebrafish Book : A Guide for the Laboratory Use of Zebrafish.

Wulliman, M.F., Rupp, B., Reichert, H., 2012. Neuroanatomy of the Zebrafish Brain: A Topological Atlas. Birkhäuser.

Yokogawa, T., Hannan, M.C., Burgess, H.A., 2012. The Dorsal Raphe Modulates Sensory Responsiveness during Arousal in Zebrafish. J. Neurosci. 32, 15205–15215.

